# MDMA-Enhanced Exposure Therapy Reverses PTSD-Like Features In a Learned Helplessness Mouse Model

**DOI:** 10.64898/2026.08.06.743271

**Authors:** Orr Shahar, Peretz Golding, Masha Chaykin, May Ben Ari, Alexander Botvinnik, Tzuri Lifschytz, Bernard Lerer

## Abstract

Post-traumatic stress disorder (PTSD) is a highly prevalent, debilitating psychiatric condition. Existing treatments are ineffective for many patients. 3,4- methylenedioxymethamphetamine (MDMA)-assisted psychotherapy has demonstrated substantial clinical efficacy but relies on prolonged, resource-intensive therapeutic protocols that limit scalability and accessibility. Here, we investigated whether combining MDMA with exposure-based intervention could enhance therapeutic efficiency in a preclinical model of PTSD-like behavior. Using a learned helplessness paradigm in mice, we identified trauma-susceptible individuals based on persistent escape failures following inescapable stress. Traumatised mice subsequently received brief treatment regimens consisting of MDMA or saline vehicle administered with or without exposure to the traumatic cue. Behavioral outcomes were tracked longitudinally using active avoidance performance as the primary endpoint, complemented by assays of anxiety-like, depressive-like, cognitive, and social behaviors. MDMA treatment markedly reduced trauma-associated behavioral deficits. MDMA combined with exposure produced rapid and sustained recovery compared to control conditions. Statistical analyses revealed significant treatment- and time-dependent effects on avoidance behavior, indicating accelerated resilience acquisition in MDMA-treated groups. Additional behavioral assays demonstrated dose-dependent effects of MDMA on anxiety- and depression-related measures. Together, these findings provide proof-of-principle that pharmacological modulation with MDMA can enhance exposure-driven behavioral recovery, supporting a strategy to integrate MDMA into more efficient and accessible PTSD treatment frameworks. This work establishes a preclinical foundation for clinical studies aimed at optimizing MDMA-assisted interventions to improve scalability and patient access.

## Introduction

Posttraumatic stress disorder (PTSD) is a severe and chronic psychiatric condition characterized by persistent fear, avoidance, and dysregulated emotional learning following exposure to trauma. Global epidemiological studies indicate that PTSD affects a substantial proportion of the population (approximately 4%) and is associated with marked functional impairment, comorbidity, and significant societal burden (1–4). Although trauma-focused psychotherapies and pharmacological treatments are considered first-line interventions for PTSD, treatment outcomes remain highly variable (5). In military populations, 60-72% of patients receiving cognitive processing therapy or prolonged exposure, retain their PTSD diagnosis post-treatment (5). Non-response rates in PTSD outcome studies can reach as high as 50% (6). Dropout rates in psychological therapies for PTSD average 16% in randomized controlled trials (7). Pharmacotherapy yields small effect sizes with dropout rates showing no significant difference from placebo (8).

PTSD involves dysregulated fear responses, including exaggerated acquisition and impaired extinction of fear memories (9). It manifests as a disorder of fear dysregulation, with abnormalities in brain circuits mediating fear and stress (10). The limitations of currently available treatments highlight a critical need to identify strategies that more effectively engage the learning processes required for durable fear reduction.

Psychedelic compounds have emerged as a promising avenue for addressing treatment-resistant psychiatric disorders. Contemporary clinical and preclinical work suggests that psychedelics can induce enduring changes in affective processing and learning while exhibiting acceptable safety profiles under controlled conditions (11–13). In the context of PTSD, a study by Catlow et al. reported that out of a number of doses of psilocybin, the lower doses (0.1, and 0.5 mg/kg) facilitated extinction of fear conditioning in mice in parallel to increased hippocampal neurogenesis (14). Among psychedelics and related compounds, the entactogen 3,4-methylenedioxymethamphetamine (MDMA) is of particular interest due to its capacity to reduce threat sensitivity while preserving emotional engagement (15). Recent syntheses of the field emphasize the relevance of psychedelic compounds for stress-related psychiatric disorders, including PTSD, and suggest that their therapeutic effects may be mediated through alterations in emotional learning and plasticity (16, 17).

Consistent with this framework, MDMA-assisted psychotherapy has demonstrated robust efficacy in multiple clinical trials for PTSD. Phase 2 and Phase 3 studies conducted by the Multidisciplinary Association for Psychedelic Studies (MAPS) have reported large and durable reductions in symptom severity following a limited number of MDMA-assisted sessions, with effects significantly greater than psychotherapy alone (18–21). While these findings represent a major advance, the therapeutic model involves extensive preparatory and integrative psychotherapy, requiring substantial time and clinical resources. As a result, scalability, cost, and patient throughput remain significant challenges for broad implementation (22). Two recent Phase 3 trials confirmed large effect sizes for MDMA-assisted therapy in moderate-to-severe PTSD, yet regulatory approval remains pending following FDA feedback in 2024 citing needs for additional safety and efficacy data in new trials. To address these barriers, emerging clinical protocols are now testing MDMA combined with massed (intensive, daily) prolonged exposure therapy (the METEMP protocol), aiming to shorten treatment duration while preserving efficacy (23). A more cost-effective therapeutic approach could be of significant value to the field

The learned helplessness paradigm, originally developed by Seligman and colleagues to investigate the effects of uncontrollable aversive events, involves exposure to inescapable electric shocks, leading to subsequent deficits in escape responding in escapable situations (24, 25). While classically employed as a preclinical model for major depressive disorder due to its induction of passive behavior and motivational deficits (26), learned helplessness captures core features of posttraumatic stress disorder (PTSD), including persistent behavioral helplessness following uncontrollable trauma, and re-experiencing symptoms triggered by trauma-associated cues in the same context (27, 28).

In humans, only a subset of individuals exposed to the same traumatic event develop PTSD, with conditional risk varying by trauma type but averaging around 4% overall and higher for interpersonal violence (2). The conditional risk of developing PTSD increases markedly with trauma severity and context; for example, the risk is approximately 3.5% following motor vehicle accidents but rising to as high as 19% following rape and other severe forms of interpersonal violence (29). In modern war situations the prevalence escalates dramatically, with lifetime PTSD reaching 15–29% among veterans of recent conflicts (30). Our modified protocol implements the classic inescapable shock induction over two days, followed by screening in weekly escapable shock sessions (30 trials), where only 30-40% of mice exhibit strong, persistent high failure rates (above 70%) lasting over 3 months, while the remainder rapidly return to low failure rates (0-10%) by week 2. This individual variability mirrors human PTSD susceptibility and supports the model’s face validity for PTSD-like indications. Here, failure percentage in escapable trials serves as a reliable index of PTSD-like behavior, with reductions reflecting fear extinction.

At a mechanistic level, exposure-based therapies are thought to operate through fear extinction and inhibitory learning processes that depend on coordinated activity within amygdala–prefrontal networks (31, 32). However, avoidance and excessive fear responding can limit engagement with trauma-related cues, thereby constraining extinction learning. Preclinical studies indicate that MDMA can facilitate fear extinction, enhance emotional learning, and modulate neural circuits implicated in threat processing (17, 33–35). Notably, most experimental studies examining psychedelic or MDMA effects in PTSD-relevant models have assessed drug effects in isolation, rather than administering the drug acutely during explicit trauma cue exposure or fear extinction sessions. Here, we investigate whether co-treatment with MDMA and exposure alleviates PTSD-like behavioral phenotypes in a mouse model, providing a proof-of-principle demonstration that pharmacological modulation of emotional state can enhance exposure-driven fear reduction and inform the development of more efficient and accessible therapeutic strategies.

## Results

### Modified learned helplessness induction yields persistent escape deficits in a subset of mice

We have established a modified active avoidance-learned helplessness model with persistent, long-lasting (up to 3 months) escape failure behaviour (Supplementary Figure 1). Our experimental protocol included pretest, inductions, tests, exposure treatment sessions, and a behavioural battery (Fig. 1).

**Figure 1.**
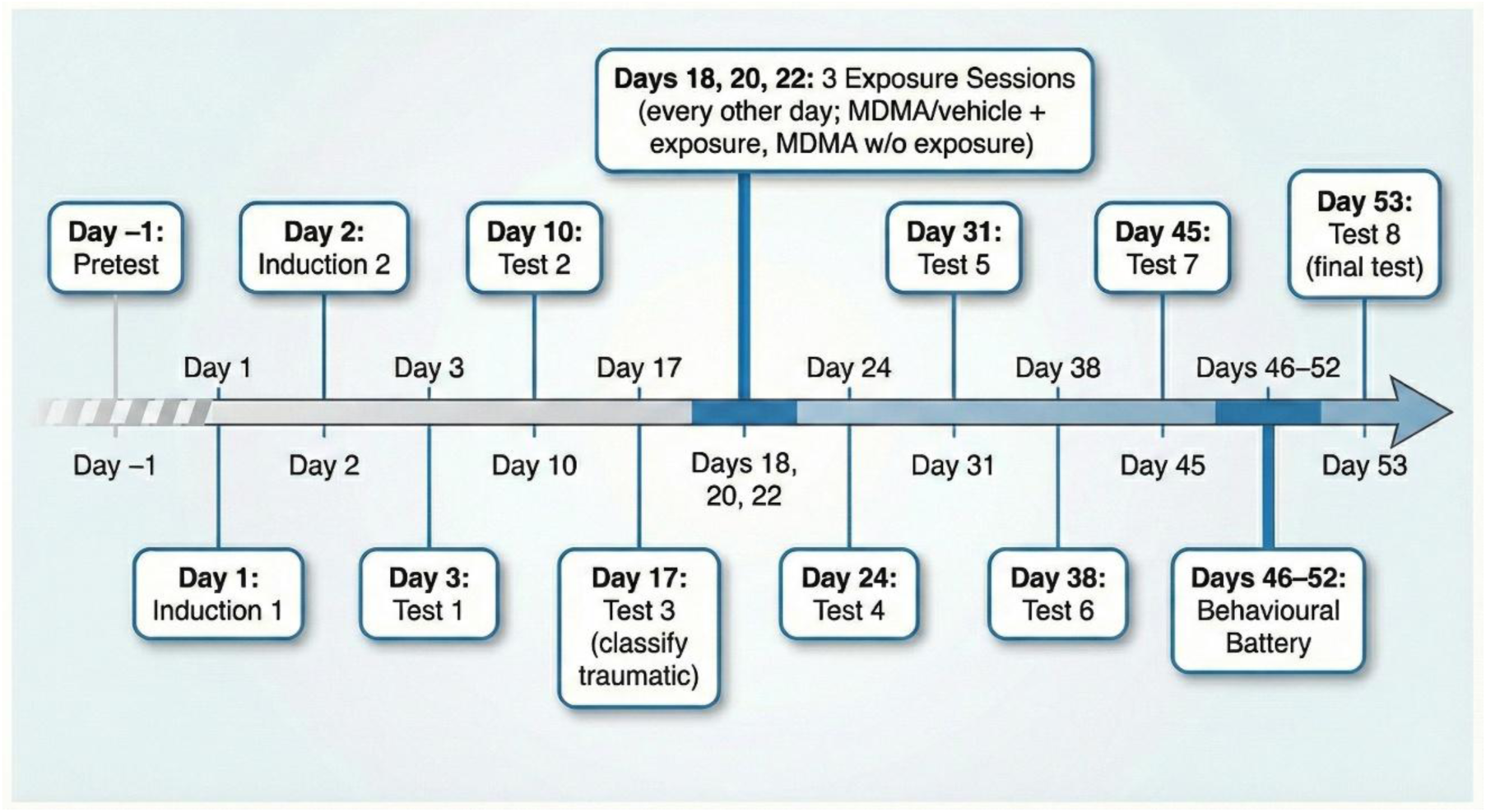
Timeline indicating different parts of the experiment and on which days they were performed.

Pretest (escapable 30 shock test) confirmed escape competency, allowing exclusion of outliers that failed more then 50% of 30 escapes. Following two consecutive days of inescapable shock induction, weekly escapable shock tests (Tests 1–3) revealed bimodal outcomes: 38% of mice (n=21) developed persistently high failure rates (>70% by Test 3) and were classified as “traumatic” (PTSD-like), while “resilient” mice (<33% failure) rapidly normalized (Fig. 2). This variability persisted without intervention.

**Figure 2.**
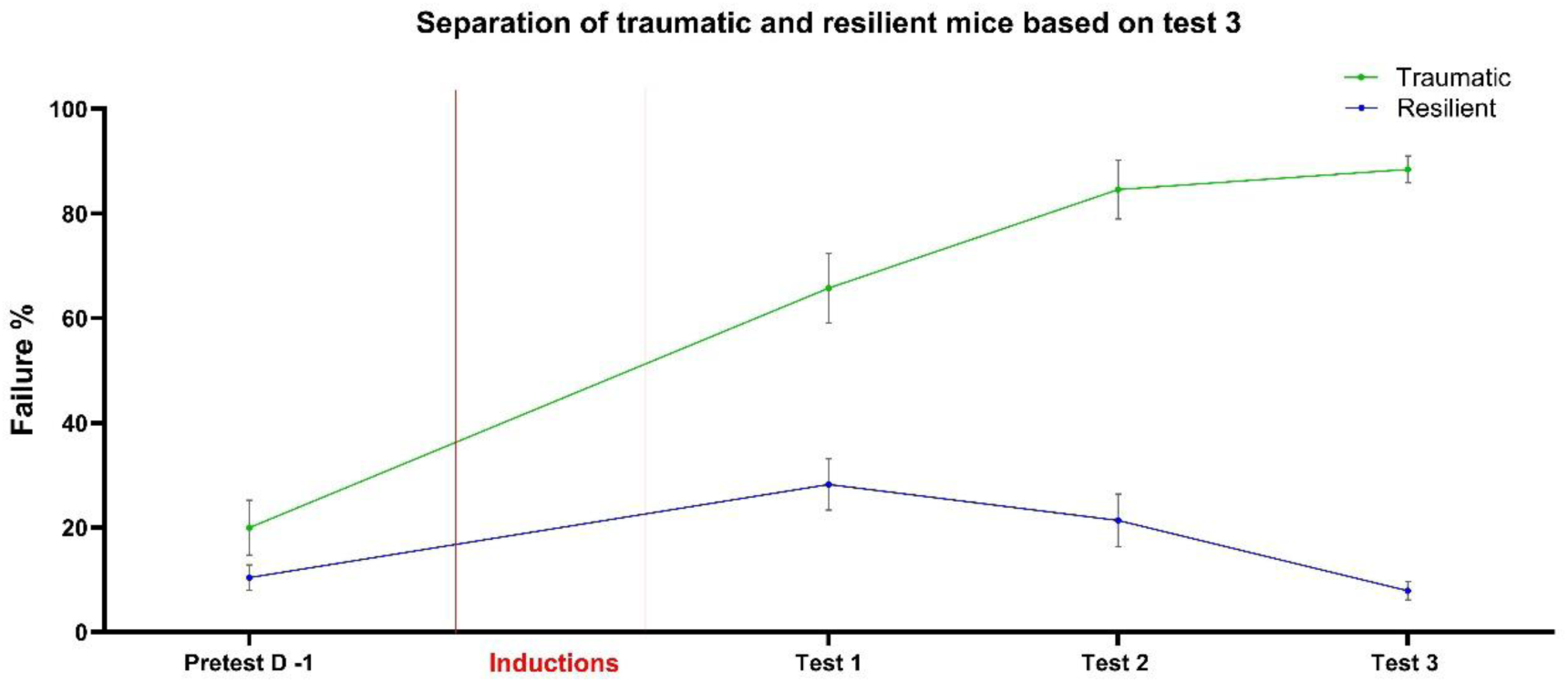
Screening of mice based on test 3 results. Mice that scored under 33% failure rate were classified as Resilient (n=33), and mice who scored above 70% failure were classified as Traumatic (n=21). Failure % is based on failure to escape shock out of 30 trials. Error bars represent SEM.

### Effect of MDMA with or without exposure on escape failures in “traumatic” mice

Traumatic mice underwent either three (every other day) MDMA 7.5 mg/kg treatments or three spaced exposure sessions (every other day) under MDMA (7.5 and 15 mg/kg) or saline vehicle between Tests 3 and 4. Two-way ANOVA revealed significant Treatment (p = 0.0001) and Treatment × Time (p < 0.0001) effects on failure rates. MDMA 7.5 mg/kg + exposure and MDMA 15 mg/kg + exposure groups achieved immediate resilience (<33% failure) from Test 4 onward, sustained through Test 8. MDMA 7.5 mg/kg without exposure showed gradual reduction, reaching resilience by Test 6 and sustaining thereafter. Vehicle + exposure maintained high failures (>70%) throughout (Fig. 3).

**Figure 3.**
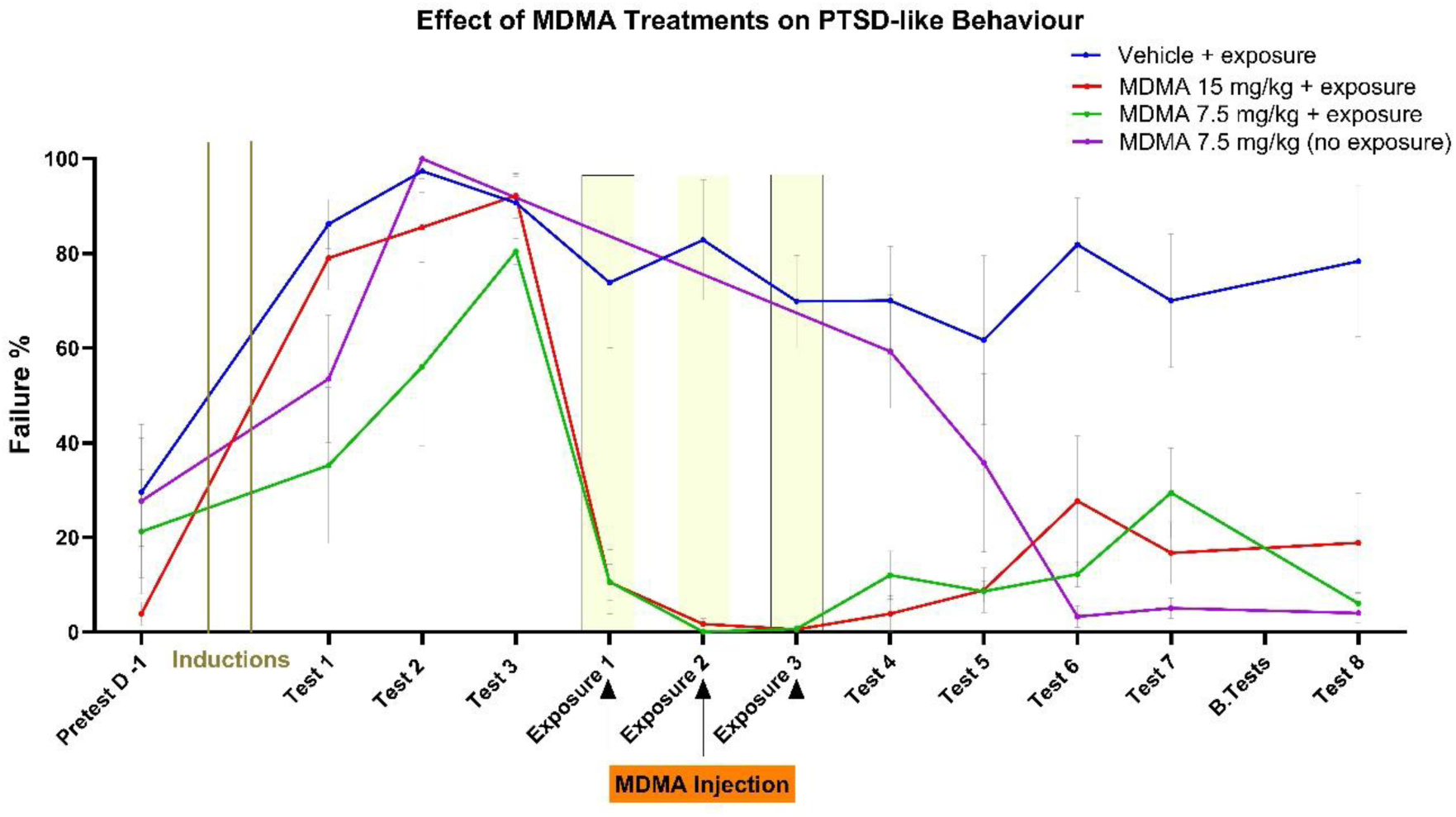
MDMA treatments reduces failure to escape stimulus shock. Failures shown are a % out of 30 trials on presented shock. MDMA treatments significantly reduced failure rates, while Vehicle stayed consistently high. Failure % is based on failure to escape shock out of 30 trials. Error bars represent SEM. (n=4-7)

### MDMA effects on anxiety-like behaviors

On day 46 mice performed the elevated plus maze (EPM) and the light/dark box. All the mice were exposed to the exposure sessions, except for the Resilient group and the MDMA 7.5 mg/kg (no exposure) group. MDMA 15 mg/kg reduced open-arm time vs. Vehicle (p = 0.0401), indicating increased anxiety-like behavior (Fig. 4). In the light/dark box, MDMA 7.5 mg/kg increased light-side time vs. Vehicle (p=0.0391), and MDMA 15 mg/kg (p = 0.0167), indicating reduced anxiety-like behavior for MDMA 7.5 mg/kg(Fig. 5). Data for both anxiety tests were analyzed by two-tailed Mann–Whitney tests with Holm–Šídák correction for multiple comparisons.

**Figure 4.**
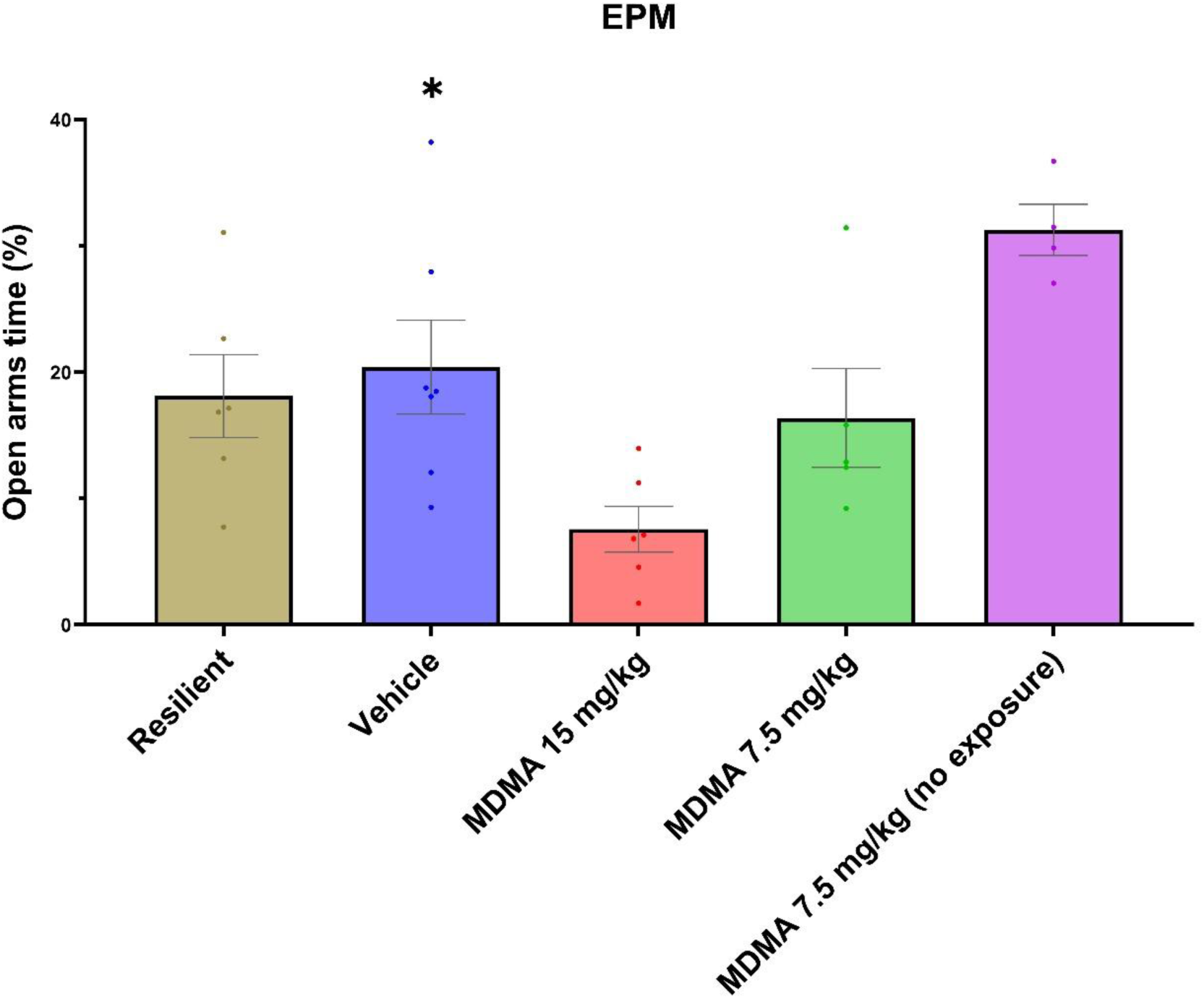
Effects of MDMA treatments on % of time spent in open arms in the EPM. MDMA 15 mg/kg reduced the time spent in open arms when compared to Vehicle. *p < 0.05 by two-tailed Mann-Whitney tests with Holm-Šídák correction for multiple comparisons. Error bars represent SEM. (n=4-7)

**Figure 5.**
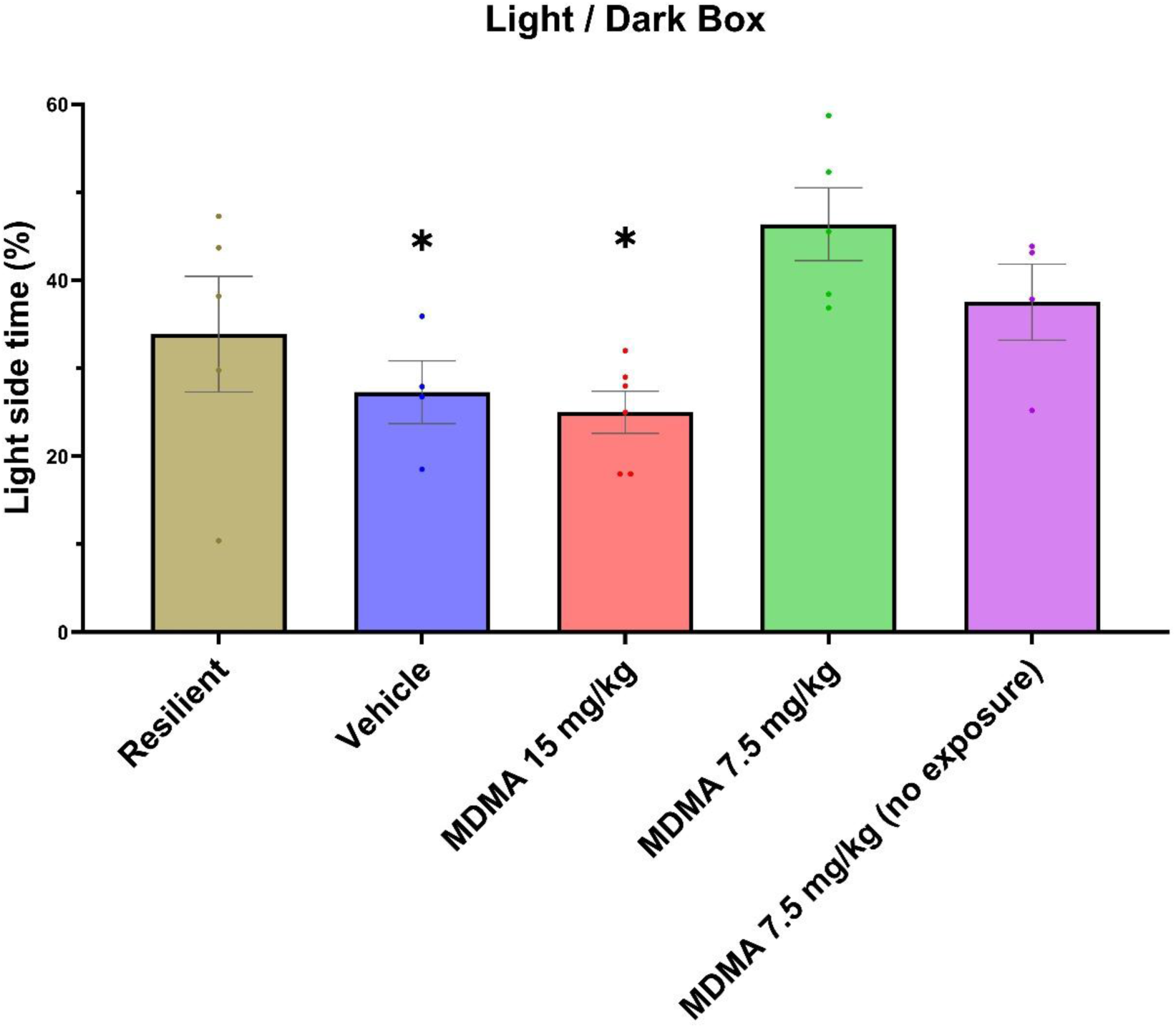
Effects of MDMA treatments on % of time spent in open arms in the light/dark box test. MDMA 7.5 mg/kg increased the time spent in light side of the arena when compared to Vehicle, and MDMA 15 mg/kg. *p < 0.05 by two-tailed Mann-Whitney tests with Holm-Šídák correction for multiple comparisons. Error bars represent SEM. (n=4-7)

### MDMA effects on depressive-like behavior

On day 47, the mice performed the forced swim test (FST). All the mice underwent to the exposure sessions, except for the Resilient group and the MDMA 7.5 mg/kg (no exposure) group. Data were analyzed by two-tailed Mann–Whitney tests with Holm– Šídák correction for multiple comparisons. MDMA 7.5 mg/kg reduced immobility vs. Vehicle (p = 0.0313), and MDMA 15 mg/kg (p = 0.0214), indicating anti-depressive-like effects (Fig. 6).

**Figure 6.**
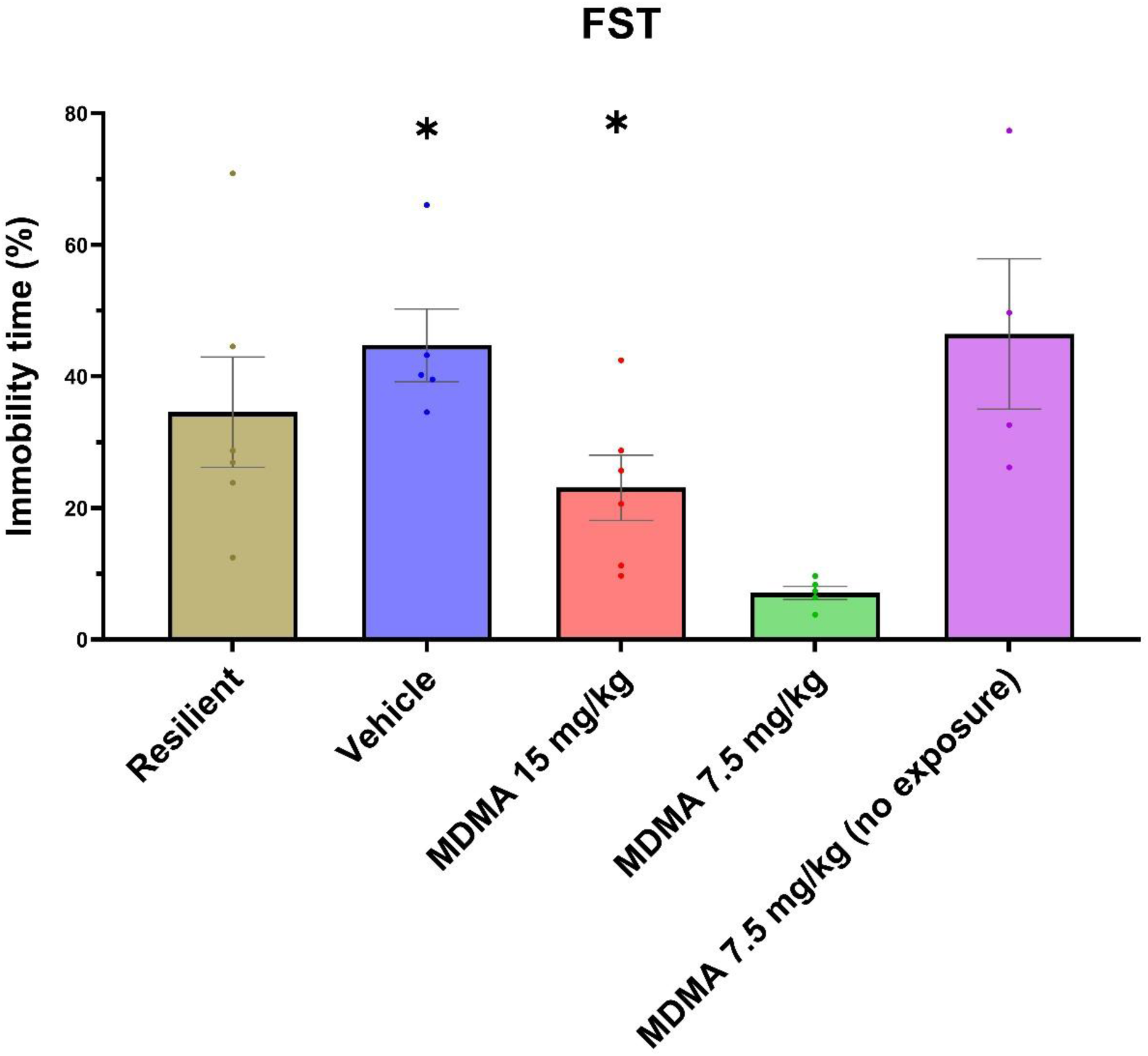
Effects of MDMA treatments on % of immobility time in the FST. MDMA 7.5 mg/kg reduced immobility time when compared to Vehicle, and MDMA 15 mg/kg. * vs MDMA 7.5 mg/kg, *p < 0.05 by two- tailed Mann-Whitney tests with Holm-Šídák correction for multiple comparisons. Error bars represent SEM. (n=4-7)

### No MDMA effects on cognition or sociability

On day 48 and 49, mice performed the T-maze, and no group differences in novel arm preference was observed (Supplementary Fig. 2). All the mice were exposed to the exposure sessions, except for the Resilient group and the MDMA 7.5 mg/kg (no exposure) group. Data were analyzed by two-tailed Mann–Whitney tests with Holm– Šídák correction for multiple comparisons. On day 52 mice performed the three-chamber test, and no differences was observed in sociability index. (Supplementary Fig. 3).

## Discussion

Our modified learned helplessness model, which incorporated susceptibility screening, effectively demonstrated PTSD-like persistent escape deficits in 36% of mice, aligning with human trauma responses where only a subset of exposed individuals develops PTSD (2). MDMA-facilitated exposure produced rapid and sustained fear extinction, with 7.5 mg/kg + exposure emerging as the optimal regimen. This dose yielded near complete and long-lasting reductions in escape failure rates without the anxiety elevation observed at 15 mg/kg + exposure.

Strengths of this study include the model’s high face and construct validity through individual variability and long-term persistence (>3 months), enabling evaluation of combined pharmacotherapy-behavioral interventions. A weakness is the small group sizes (n=4–7), which limits statistical power for subtle effects, though significant outcomes were robust.

An interesting observation was that MDMA 7.5 mg/kg administered during trauma-cue exposure reduced anxiety-like behavior in the light/dark box (increased time in the light compartment, Fig 5) and depressive-like behavior in the forced swim test (decreased immobility, Fig 4) compared with vehicle, whereas the same dose given without exposure did not. This suggests that pairing MDMA with an aversive context may confer additional affective benefit, possibly by promoting resilience or attenuating avoidance of mildly aversive stimuli. These results converge with and extend recent preclinical work showing that MDMA is most effective when administered in conjunction with trauma-cue reactivation (36). While Arluk and colleagues demonstrated this principle in rats using a single brief memory reactivation session in a predator-scent stress model, the present study is the first to show in mice that repeated MDMA-enhanced exposure sessions produce rapid, robust, and sustained reversal of persistent PTSD-like escape deficits in a learned-helplessness paradigm with high face validity for individual susceptibility and chronicity. Our findings therefore provide direct preclinical proof-of-concept that MDMA can be used to enhance the efficiency of exposure-based interventions, supporting the development of more scalable clinical protocols.

In addition, our results converge with recent clinical work on MDMA-enhanced massed exposure. In a case study from the METEMP protocol, a 57-year-old woman with severe PTSD (baseline CAPS-5 = 85) achieved full remission (CAPS-5 = 1) one week and one month after a 10-day massed prolonged exposure course paired with a single MDMA dose (23).

These findings imply that MDMA-assisted exposure could accelerate PTSD remission in resistant cases, integrating with established therapies like prolonged exposure (37). Given MDMA’s clinical safety profile, minimal neurotoxicity at therapeutic doses, and low adverse events in Phase 3 trials (18, 38), our proof-of-principle supports advancing to human trials. These preclinical findings provide timely translational support for the expanding clinical evaluation of MDMA-enhanced massed exposure approaches currently underway, including the Emory METEMP open-label pilot (NCT05746572) and the DoD-funded randomized controlled trial of MDMA-assisted massed PE in ∼100 military participants (39, 40). This could transform PTSD treatment, reducing societal burdens like chronic disability and suicide risk, offering hope for trauma survivors worldwide by enabling efficient, accessible recovery.

## Methods

### Animals and housing

Male C57BL/6J mice (n=60, aged 11 weeks at experiment start; Jackson Laboratory) were housed in groups of 4–5 per cage under standard conditions (ad libitum food and water, 12-h light/dark cycle, 22°C). Animals were acclimated for 1 week before the pretest. Experiments were conducted in accordance with AAALAC guidelines and were approved by the Authority for Biological and Biomedical Models, Hebrew University of Jerusalem, Israel, Animal Care and Use Committee protocol number MD-24-17534-4. All efforts were made to minimize animal suffering and the number of animals used.

### Active avoidance apparatus

Experiments used the Hugo Basile Active Avoidance set-up for Mice and Rats (Shuttle-Box; Product Codes 40532/40533), comprising two arenas separated by a wall with a small hole allowing easy passage. The floor consisted of metal bars for programmable shock delivery (0.3 mA).

### Learned helplessness induction and testing protocols

One day before induction, a pretest (30 escapable shocks: 0.3 mA, 10 s duration, 30 s intervals) assessed baseline escape ability; mice with high failure rates (outliers from cage mates) were excluded. Induction involved two consecutive days of 200 inescapable shocks (0.3 mA, 2 s duration, 10–30 s random intervals). Weekly tests (Tests 1–3, 4–8) comprised 30 escapable shocks (0.3 mA, 10 s duration, 30 s intervals). After Test 3, mice were classified as "traumatic" (PTSD-like; >70% failure rate; n=22, 36% of cohort) or "resilient" (<33% failure rate). Traumatic mice were randomized into four groups: Vehicle + exposure (n=7), MDMA 7.5 mg/kg + exposure (n=5), MDMA 7.5 mg/kg without exposure (n=4), MDMA 15 mg/kg + exposure (n=6). A random subset of resilient mice served as a comparator for behavioral tests.

### MDMA administration and exposure sessions

d,l-MDMA·HCl (98.5% purity by HPLC, 81.9% freebase; Lipomed AG, Arlesheim, Switzerland) or Vehicle (0.9% saline) was administered intraperitoneally (i.p.) 30 min before sessions. Exposure therapy occurred in the week between Tests 3 and 4, with three sessions every other day (e.g., Sunday, Tuesday, Thursday) in the escapable shock paradigm (30 trials). MDMA without exposure groups received injections on the same schedule but remained in home cages. Weekly testing resumed post-exposure (Tests 4–8; see Fig. 1 for timeline).

### Behavioral test battery

One week after Test 7, mice underwent a behavioral battery. Elevated plus maze (EPM): 5-min free exploration in a plus-shaped maze, two arms “open” without walls, and two arms “closed” with closed walls (arms: 30 × 5 cm, elevated 50 cm); time in open arms measured anxiety-like behavior. Forced swim test (FST): 6-min immersion in a cylinder (25 cm diameter, 30 cm water depth, 25°C); immobility time throughout assessed depressive-like behavior. Three-chamber sociability test: 10-min habituation followed by 2 x 10-min trials with a stimulus mouse (familiar) vs. empty cup, then novel vs. familiar mouse; time in chambers and near cups quantified social preference. T-maze: 5-min habituation with one arm of the T closed, then a day after, 5-min exploration with all arms open; preference for novel arm assessed spatial memory. Light/dark box: 5-min exploration in a box (40 × 20 cm) divided into light (illuminated) and dark compartments; time in light side measured anxiety-like behavior.

### Statistical analysis

Data are presented as mean ± SEM. Failure rates were quantified as the number of failed escapes attempts out of 30 shocks per test session. Two-way ANOVA was performed on the active avoidance test. Owing to small sample sizes in the behaviour tests (n = 4–7 per group), formal normality tests were underpowered, and one-way ANOVA could not be reliably performed. Data distributions were therefore assessed by visual inspection of histograms and Q–Q plots, which showed no clear deviations from normality. The primary hypotheses were defined a priori (comparison of each MDMA treatment group versus vehicle + exposure, and MDMA 15 mg/kg versus MDMA 7.5 mg/kg). Behavioral data were analyzed using two-tailed Mann-Whitney tests for these planned comparisons, with Holm–Šídák correction for multiple testing to control the family-wise error rate. Statistical significance was set at p < 0.05. All analyses were performed using GraphPad Prism (v9).

Acknowledgement of Generative AI and AI-assisted Technologies Used in Writing Grok (xAI) was used to assist with editing selected sections of the manuscript, specifically to improve clarity, flow, and overall reader experience. Google Gemini was used to help create Figure 1.

## Data Availability

The data that support the findings of this study are available from the corresponding author upon reasonable request.

## Supporting information

Shahar-Supplementary Information

## Acknowledgements

The research was supported in part by Negeb Labs. O.S. is supported by the President of the State of Israel Scholarship for Excellence and Scientific Innovation, and the Hadassah BrainLabs Center for Psychedelic Research. We thank the staff of the Hebrew University Animal Facility for excellent animal care.

## Author contributions

O.S. designed and performed all experiments, analyzed data, prepared figures and wrote the manuscript. P.G., M.C., and M.B.A. assisted with behavioral experiments and data collection. A.B. assisted with statistical analysis and manuscript editing. T.L. and B.L. conceived and supervised the project, secured funding, and edited the manuscript. All authors reviewed and approved the final manuscript.

## Competing interests

B.L is a consultant to Negev Labs.

## Supplementary materials

Supplementary materials are provided.

## Notes

### Competing Interest Statement

Bernard Lerer is a Consultant to Negev Labs

