## Supplementary material for "MDMA-Enhanced Exposure Therapy Reverses PTSD-Like Features In a Learned Helplessness Mouse Model": Shahar-Supplementary Information

<u>Supplementary Data for MDMA-Enhanced Exposure Therapy Reverses PTSD-Like Symptoms in a Preclinical Learned Helplessness Mouse Model</u>

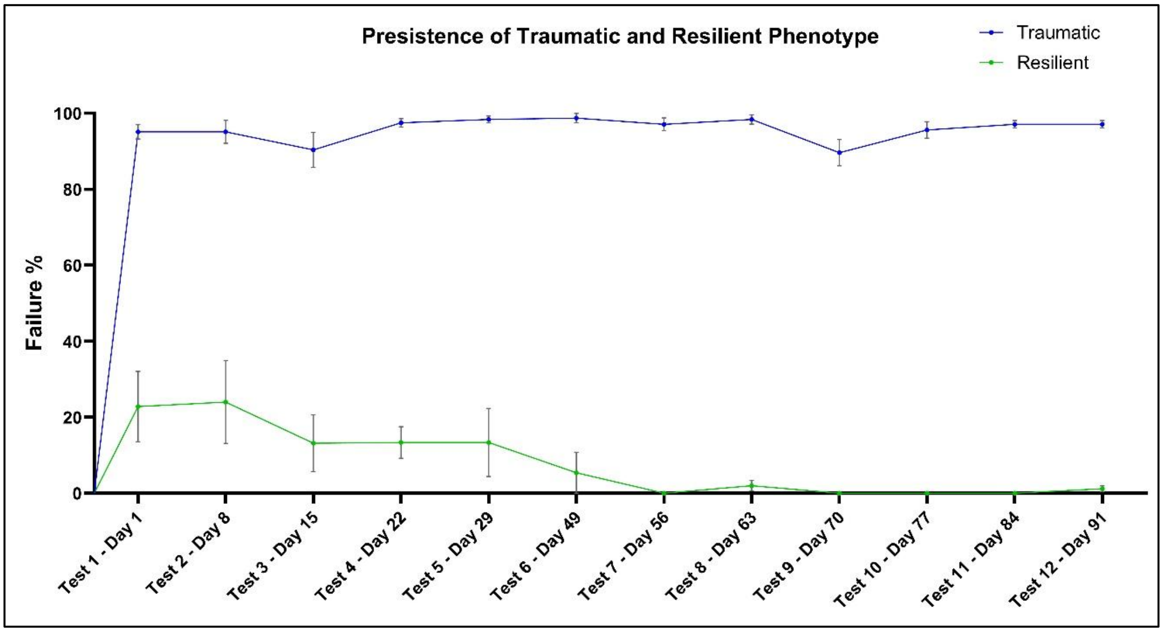


*Supplementary Figure 1.* Screening of mice based on test 3 results. Mice who scored under 33% failure rate were classified being resilient, and mice who scored above 70% failure were classified traumatic. Failure % is based on failure to escape shock out of 30 trials. Error bars represent SEM. (n=5-8)

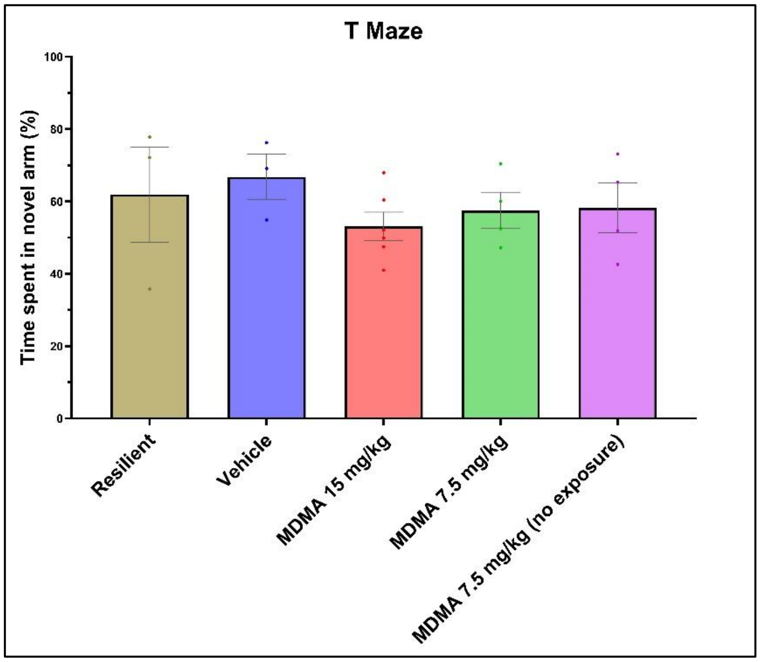


*Supplementary Figure 2.* Effects of MDMA treatments on time spent in novel arm in the T maze test. No difference was observed between the groups by two-tailed Mann-Whitney tests with Holm-Šídák correction for multiple comparisons. Error bars represent SEM. (n=4-7)

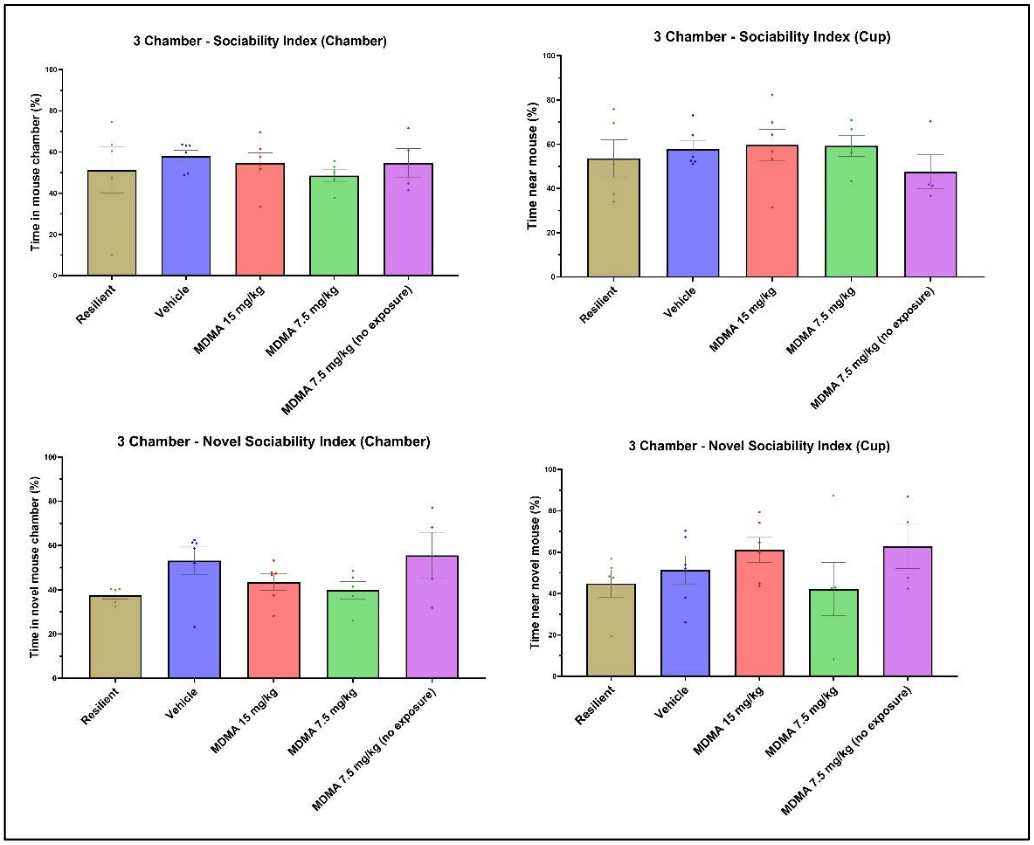


*Supplementary Figure 3.* Effects of MDMA treatments on time spent around social arousal the 3-chamber test. No difference was observed between the groups by two-tailed Mann-Whitney tests with Holm-Šídák correction for multiple comparisons. Error bars represent SEM. (n=4-7)
